# Lower GABA levels in the posterior cingulate are linked with poorer episodic memory in healthy older adults

**DOI:** 10.1101/2025.03.18.643988

**Authors:** Poortata Lalwani, Casey Vanderlip, Craig Stark

**Affiliations:** University of California, Irvine

## Abstract

Age-related deficits in episodic memory and mnemonic discrimination are associated with an increased risk of neurodegenerative diseases, such as Alzheimer’s disease (AD) (Stark et al., 2013). While much research has focused on hippocampal contributions to these age-related changes (Stark et al., 2019), less is known about the role of posterior cingulate cortex (PCC) especially reduced inhibition in episodic memory deficit. PCC has connections to the medial temporal lobe and is linked to memory declines (Greicius et al., 2004). It is also one of the most vulnerable regions to amyloid deposition in AD (Yokoi et al., 2018). This study hypothesized and found that age-related declines in GABAergic function (brain’s major inhibitory neurotransmitter) within the PCC contributes to individual differences in memory performance in healthy older adults. Using Magnetic Resonance Spectroscopy, we measured GABA levels in the PCC in 22 healthy younger and 30 older adults. We assessed episodic memory using Rey Delayed Auditory Verbal Learning Test (RAVLT) and Mnemonic Similarity Task (MST). We found that both raw GABA levels and episodic memory performance are lower in older adults compared to young. This reduction in GABA levels is subserved by age-related changes in tissue-composition as evidenced by no age-group differences in corrected GABA levels. More importantly, lower GABA levels (independent of tissue-correction) were associated with poorer episodic performance including delayed recall and mnemonic discrimination. This research suggests that therapeutically targeting posterior cingulate GABA levels might help slow or alleviate memory decline.

**Significance Statement:** This study provides novel insights into the role of posterior cingulate cortex (PCC) GABA+ levels in age-related memory deficits. Our findings demonstrate that lower PCC GABA+ levels in older adults are associated with poorer performance on episodic memory tasks, particularly those involving mnemonic discrimination and word-list learning. This research expands on the growing body of literature linking GABAergic dysfunction to age-related cognitive impairments and suggests that GABAergic changes in the PCC contribute to episodic memory deficits. Importantly, our results highlight the potential of targeting PCC GABA levels as a therapeutic strategy to slow or mitigate memory decline in aging. These findings also offer promising avenues for future research into early biomarkers for Alzheimer’s disease and other neurodegenerative conditions.

## Introduction

Episodic memory declines early in healthy aging with deficits emerging as early as the third decade of life (Cansino, 2009). This is, in part, due to deficits in mnemonic discrimination, which is the ability to distinguish between similar events (Yassa and Stark, 2011). Mnemonic discrimination critically supports episodic memory, and this ability declines drastically with age (Stark et al., 2013). More importantly, there is significant interindividual variability in both episodic memory and mnemonic discrimination within older adults (Stark et al., 2013; Tromp et al., 2015). Importantly, older adults with deficits in episodic memory are at high risk for future cognitive decline and neurodegenerative disorders, such as Alzheimer’s disease (Bäckman et al., 2001; Lee et al., 2018; Vanderlip et al., 2024b).Therefore, it is critical to identify the mechanisms that contribute to the individual variability in episodic memory within healthy older adults.

While significant work has investigated the role of hippocampus and adjacent structures in episodic memory, less work has focused on the age-related differences in cortical regions with hippocampal projections and its role in episodic memory declines. However, cortical regions with strong projections to the hippocampus, such as the posterior cingulate cortex, play a critical role in memory capacity. Specifically, decreased volume and microstructural changes within the posterior cingulate, which has strong projections to the medial temporal lobe, has been linked to age related deficits in episodic memory (Chua et al., 2009; Stanislav et al., 2013; Gefen et al., 2015). Importantly, the posterior cingulate is also one of the most vulnerable regions to amyloid deposition in Alzheimer’s disease suggesting that alterations in this region may play a role in the drastic declines in episodic memory seen in this condition (Mattsson et al., 2019). It is unclear whether neurochemical alterations specifically, reduced inhibition within posterior cingulate, relates to memory deficits in cognitively healthy older adults.

Gamma-aminobutyric acid (GABA) is the most widespread inhibitory neurotransmitter in the brain and is responsible for modulating synaptic transmission, synchronizing cortical network activity, and promoting neuronal development and relaxation (Ngo and Vo, 2019). However, GABAergic function declines in healthy aging. A study in a large, healthy, older population indicates that GABA concentrations in both frontal and posterior regions decrease with age (Porges et al., 2017). Similarly, a cross-sectional study of healthy adults aged 20–76 years found that GABA concentrations decrease with age after adolescence in both the frontal region and parietal regions (Gao et al., 2013). Thus, we hypothesized an age-related reduction in GABA concentrations in the posterior cingulate cortex.

Additionally, GABA levels have been associated with a wide range of cognitive and memory tasks (Makkar et al., 2010; Heaney and Kinney, 2016). Higher GABA concentrations predicts better global cognitive performance (evaluated using the MoCA) in healthy older adults (Porges et al., 2017). Similarly, within the frontal cortex, higher GABA concentrations in the dorsolateral prefrontal cortex (DLPFC) correlates with better working memory under high load task conditions (Yoon et al., 2016; Ragland et al., 2020). In the hippocampus, higher baseline GABA levels have been associated with better retrieval performance in an associative learning paradigm (Spurny et al., 2020). Based on the role of posterior cingulate in episodic memory (Dennis and Peterson, 2012; Leech and Sharp, 2014; Papma et al., 2017; Natu et al., 2019; Edde et al., 2020), we hypothesized that reduced GABA levels in the posterior cingulate might predict individual differences in episodic memory and mnemonic discrimination within healthy older adults.

To test this hypothesis, we used Magnetic Resonance Spectroscopy to measure GABA levels in the posterior cingulate cortex and assessed episodic memory using the Rey Delayed Auditory Verbal Learning Test (RAVLT) and mnemonic discrimination using the Mnemonic Similarity Task (MST). We chose these tasks given that performance on both tasks declines rapidly with age and lower performance on these tasks is associated with increased risk of AD pathology (Estévez-González et al., 2003; Ricci et al., 2012; Stark et al., 2013; Vanderlip et al., 2024c).

## Methods

### Participants

We analyzed the data from 22 young (age 18-29 years) and 30 older (age 65 and above) adults who completed both posterior-cingulate GABA imaging session and behavior testing. All participants were recruited from Irvine and the surrounding area, were right-handed, native English speakers, and had normal or corrected to normal vision. All participants scored 23 or more on the Montreal Cognitive Assessment (MOCA). All the sessions described below took place at the University of California’s Functional MRI Laboratory, Irvine, California (See **Table 1** for more demographic details). The following were subject inclusion and exclusion criteria:

**Table 1.**

|  | YA | OA |
| --- | --- | --- |
| n | 22 | 30 |
| Age (std) | 28.45 (4.74) | 76.03 (6.53) |
| Female (%) | 12 (54.5) | 16 (53.3) |
| Education (std) | 15.64 (2.67) | 17.07 (2.20) |
| Race |  |  |
| White (%) | 11 (50) | 27 (90) |
| Asian (%) | 7 (31.8) | 3 (10) |
| Black (%) | 2 (9.1) | n/a |
| Multiple (%) | 2 (9.1) | n/a |
| **Demographics** |  |  |

#### Study Inclusion Criteria

Age 18–29 years or 65 years and older, Right-handed, Native English speaker, Healthy (i.e., no known debilitating conditions, mental illness, or head trauma), MOCA>23

#### Study Exclusion Criteria

Glaucoma, Respiratory problems, Past or present chemotherapy, Immune system disorder, Kidney or liver disease, Motor control problems, Psychotropic medication, Current depression or anxiety, or occurrence of depression/anxiety within 5 years,

Concussion with unconsciousness for 5 min or more, Pregnancy or attempting to become pregnant, More than 4 alcoholic drinks per week for women, more than 6 for men, History of drug or alcohol abuse or addiction, Weight greater than 250 pounds, MRI incompatibility (claustrophobic, foreign metallic objects, pacemaker, etc.), Currently taking drugs known to interact with GABA levels like GABApentin, estrogen therapy, pregabalin etc.

### Behavioral Testing

#### Rey Auditory Verbal Learning Task (RAVLT)

The RAVLT, a gold-standard neuropsychological assessment, was used to assess episodic memory. Participants were read a list of 15 words (List A) and recalled as many as possible across five learning trials. A second list of 15 words (List B) was then presented for interference, followed by immediate recall of List A. After a 20-minute delay, delayed recall of List A was assessed, along with recognition memory using a yes/no task that included distractors. Our primary outcome measure was delayed recall of list A (score out of 15).

#### Mnemonic Similarity Task (MST)

The MST is a mnemonic discrimination task that critically relies on the hippocampus and associated regions (Kirwan and Stark, 2007; Stark et al., 2019). During the encoding phase, participants viewed a series of images of everyday objects and identify whether each item is more likely to be found indoors or outdoors. In the subsequent recognition phase, participants viewed a new series of images, which included exact repetitions (targets), items that are similar, but not identical (lures), and completely new images (foils). Participants respond to each image as “old,” “similar,” or “new.” A recognition memory score (REC) is calculated as proportion of targets accurately identified as “old”, minus foils falsely characterized as “old” (i.e. hits minus false alarms). Mnemonic discrimination was assessed using the lure discrimination index (LDI), calculated as the proportion of lures correctly identified as “similar” minus the proportion of foils incorrectly identified as “similar.” Similar to prior work, individuals with a REC score of less than 0.5 were excluded from analyses (Radhakrishnan et al., 2022; Stark et al., 2023).

#### Trail Making Test (TMT)

The TMT was administered to assess processing speed, cognitive flexibility, and executive function. In Part A, participants were required to connect numbered circles (1 to 25) in ascending order as quickly as possible. In Part B, participants alternated between numbered and lettered circles (e.g., 1 → A → 2 → B) in ascending order, requiring additional cognitive flexibility. Performance was measured by the time taken to complete each part (Trails A and Trails B). Our primary outcome measure was the time to complete Trails B, with longer completion times indicating greater difficulty with cognitive flexibility.

### Neuroimaging

MRI data were collected using a 3T Siemens Magnetom Prisma scanner with a 32-channel head coil. A high-resolution T1-weighted structural image was collected using MPRAGE vNAV motion-corrected sequence with GRAPPA PAT mode, in sagittal direction and the following parameters: TR = 2500ms; TE = 2.9 ms; TI = 1070 ms; Acceleration Factor PE= 2; Flip Angle = 8degrees; FOV = 256 mm; 176 slices with thickness = 1 mm; no spacing; interleaved; acquisition time = 7 min 12 seconds and voxel size (1 × 1 × 1 mm)

The MRS data was obtained from 3cm x 3cm x 3cm voxels placed in the left posterior cingulate region. The voxels are placed just along the midline minimizing overlap with the dura. They were rotated such that one of the edges is parallel to the cingulum boundary minimizing overlap with white matter. Note that these are large voxels, and the primary concern is to minimize overlap with non-cortical regions like ventricles, brainstem, dura, midline that affect contamination and good shim (FWHM less than 15) while maximizing the coverage of region of interest. In this case the left posterior cingulate regions were targeted but the voxel covered parts of cuneus, precuneus, superior parietal lobule and middle cingulum. We used a MEGA-PRESS sequence with the following parameters to obtain MR spectra: TE=68ms, TR=2020ms, bandwidth = 1200 Hz, spectral width=2400Hz, water suppression bandwidth = 50Hz, Frequency selective editing pulses (16ms) applied at 1.9ppm (ON) & 7.5 ppm (OFF), 240 transients (120 ON interleaved with 120 OFF) of 2080 data points. A separate water reference sequence was obtained with 12 averages, TR = 2000ms, TE = 30ms, no water suppression and bandwidth = 1340Hz. Total scan time was 8 min 45 seconds.

### GABA Quantification

We used Gannet 3.3.2, a MATLAB based toolbox, to estimate GABA+/Water levels based on the MEGA-PRESS difference spectra in each of the MRS voxels (Edden et al., 2014). All the time-domain data were phase corrected, and frequency corrected using robust spectral registration and filtered with 3-Hz exponential line broadening. The GABA levels were scaled to water and expressed in institutional units by Gannet. Gannet quantifies GABA levels by fitting a five-parameter Gaussian model to the MR spectrum between 2.19 and 3.55 ppm while the water peak is modelled using a Gaussian-Lorentzian function. The MEGA-PRESS editing scheme also results in excitation of coedited macromolecules (MM), which can contribute up to 45% to the edited signal around 3ppm overlapping with the GABA peak. Thus, all GABA values are reported as GABA+ (i.e., GABA + MM) in the present study. Gannet’s integrated voxel-to-image co-registration procedure produces a binary mask of the MRS voxel. Using an SPM-based segmentation function, Gannet estimates the tissue composition (voxel fractions containing Cerebrospinal Fluid (CSF), Grey Matter (GM) and White Matter (WM)). Gannet then estimates a tissue-corrected GABA+ value that accounts for the fraction of grey matter, white matter, and CSF in each MRS voxel as well as the differential relaxation constants and water visibility in the different tissue types. Based on a quality control check of the spectra (flagged if the value was more than 3 standard deviations from mean, the fit error was greater than 10% for water, or the fit error was greater than 15% for GABA). Total 1 young and 1 old participant was flagged and excluded from the analysis. Since these are arbitrary thresholds, during our analysis we also regressed out the fit error for water and GABA to account for estimate differences caused by fit errors.

### Statistical Analysis

Statistical analyses were conducted using R. Figures were plotted using the ggplot2 package and the lme4 package was used to perform the linear mixed effects analysis. Correlations were run using Pearson cor.test and groups were compared using Students t.test. We used a Cook’s Distance greater than 4/sample size to flag and remove outliers from each of mixed effects models reported in the results.

## Results

### Older adults have poorer memory performance

Consistent with previous research, we found that older adults have lower delayed recall scores on RAVLT test (t (46.88) = 4.95, p = 9.9e-6, MEAN_OLD_ = 9, MEAN_YOUNG_ = 13.5). Previous research has shown that the MST’s REC score is stable with age, but that it’s LDI metric declines steadily with age (Toner et al., 2009; Holden et al., 2013; Stark et al., 2013, 2015, 2023; Huffman and Stark, 2017; Stark and Stark, 2017). Here, however, even after excluding participants with less than 50% recognition on the MST (a threshold often used to identify potential non-compliance), we found that older adults had lower recognition (REC) scores than younger adults (t (45.6) = 3.29, p = 0.002, MEAN_OLD_ = 0.76, MEAN_YOUNG_ = 0.85). This effect was primarily driven by younger adults having higher than normative recognition performance on the MST (see Discussion for more details). Finally, we also found lower lure discrimination index (LDI) in older adults compared to young (t (36.1) = 3.42, p = 0.002, MEAN_OLD_ = 0.18, MEAN_YOUNG_ = 0.36). Similar to prior research (Stark and Stark, 2017; Vanderlip et al., 2024a), the recognition score in MST was not correlated with LDI within older adults (r (24) = 0.1, p = 0.6).

### GABA levels are lower in older adults

GABA/H_2_O levels measured in the posterior cingulate were lower in older adults than younger adults (t (46) = 2.09, p = 0.042, MEAN_OLD_ = 1.78, MEAN_YOUNG_ = 1.94, Figure 1C). The effect of age-group on GABA levels was reliable even when using residuals after regressing out GABA and water fit errors (t(47.4) = 2.04, p = 0.046). However, when using GABA levels estimated after accounting for tissue composition difference, younger and older adults did not reliably differ in their GABA levels (t(45.4) = −0.08, p = 0.93, Figure 1D). This suggests that age-related differences in GABA+ levels might be driven by age-related declines in grey-and white-matter volume. Nevertheless, tissue-corrected GABA+ levels were significantly correlated with GABA+ levels (Figure 1E) in both older (r(26) = 0.85, p < 0.001) and younger adults (r(19) = 0.97, p < 0.001).

**Figure 1.**
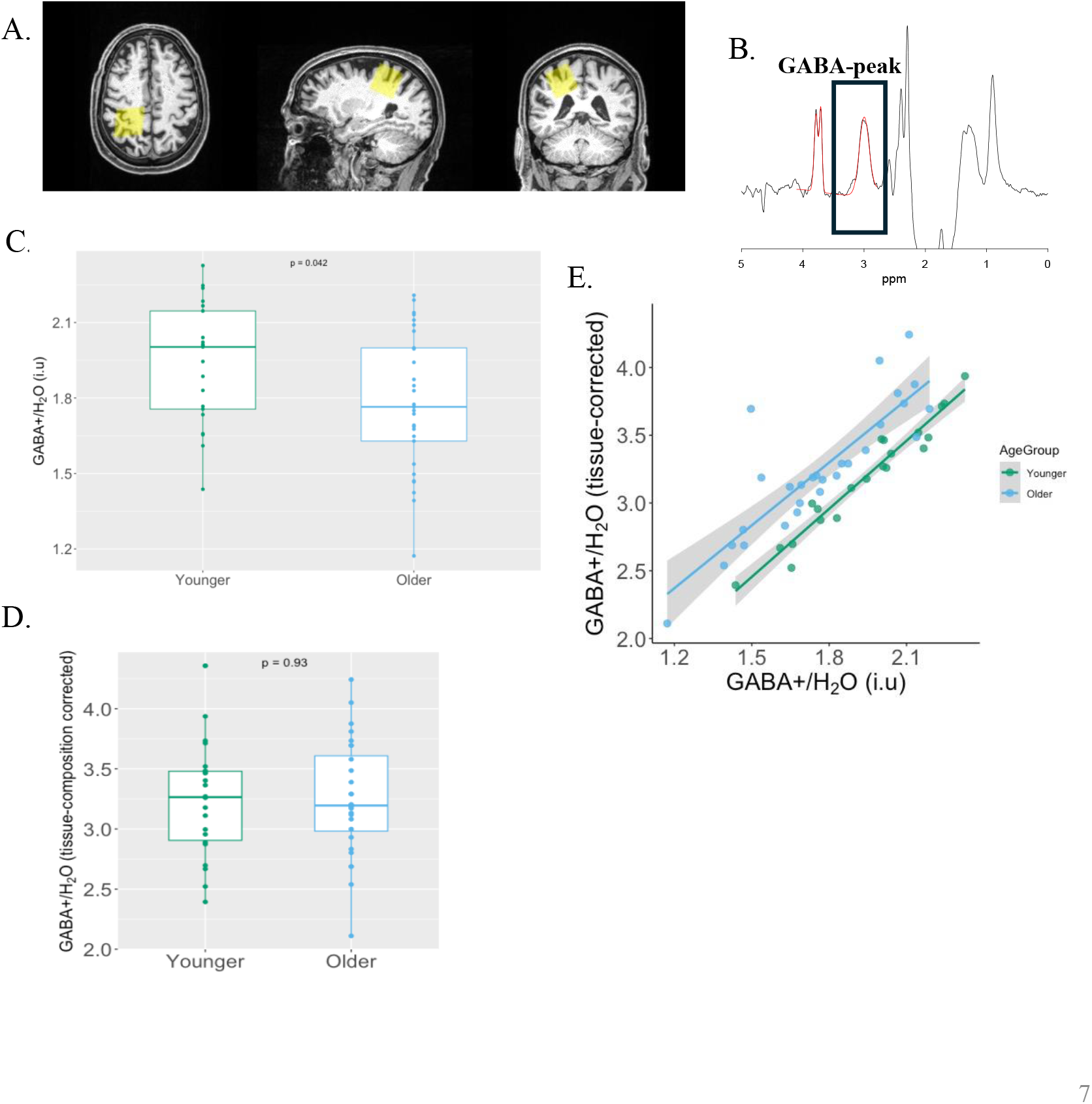
Age-related changes in GABA levels measured using MRS. A. Voxel placement example. B. Example spectra from a participant. GABA peak is at 3 ppm. C. Older adults have significantly lower raw GABA+ levels in PCC (t(46) = 2.1). D. Tissue-composition corrected GABA+ levels are not different between younger and older adults (t(45) = −0.1). E. Tissue-composition corrected GABA+ levels are strongly correlated with raw GABA+ levels within older adults (r(26) = 0.85) and within younger adults (r(19) = 0.97).

### GABA levels predict memory performance in older adults

We hypothesized that GABA+ levels in posterior cingulate would be linked with individual differences in memory performance within healthy older adults. We found that GABA+ levels were reliably correlated with delayed RAVLT (r(27) = 0.47, p = 0.008, Figure 2A) and some evidence for relationship between GABA and MST recognition (r(25) = 0.35, p = 0.07, Figure 2B) as well as MST lure discrimination (r(25) = 0.33, p = 0.09, Figure 2C) within older adults even after controlling for age, errors in GABA and water estimation. Largely, this pattern remained similar even when using tissue-composition corrected GABA+ estimates with RAVLT (r(26) = 0.27, p = 0.16, Figure 2D), MST recognition (r(25) = 0.39, p = 0.04, Figure 2E) and MST lure-discrimination (r(24) = 0.41, p = 0.03, Figure 2F). Importantly GABA+ levels are not correlated with Trails Making B performance within older adults (r (26) = −0.14, p = 0.46 (tissue-corrected: r (26) = −0.025, p = 0.9) showing domain specificity for individual differences with memory.

**Figure 2.**
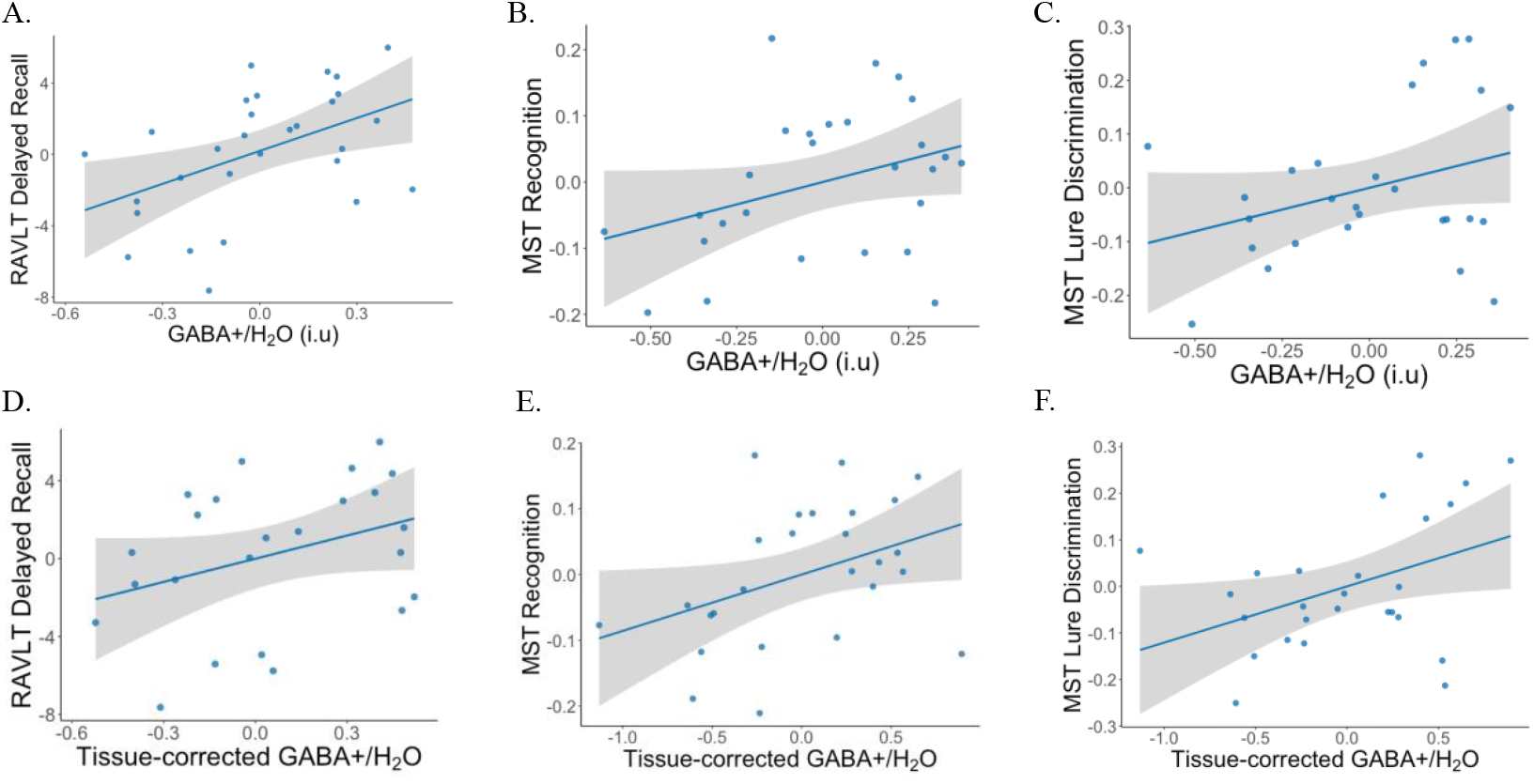
GABA levels are associated with episodic memory in older adults. A. After regressing out age, errors in GABA and water estimation, raw GABA+ levels within older adults are correlated with RAVLT delayed recall r(27) = 0.47. B. With MST recognition (r(25) = 0.35) and C. MST LDI (r(25) = 0.33) D. After regressing out age, errors in GABA and water estimation, tissue-corrected GABA+ levels within older adults show a weak relationship with RAVLT delayed ecall r(26) = 0.27, E. With MST recognition (r(25) = 0.39) and F. With MST LDI (r(24) = 0.41)

## Discussion

In this study, we replicated prior research showing age-related deficits in episodic memory, which were assessed here using the mnemonic discrimination task (MST) and the word-list learning task (Rey Auditory Verbal Learning Test, RAVLT). Moreover, we also found a novel association between lower posterior cingulate GABA+ levels and poorer memory performance in older adults. These results are discussed below in the context of existing literature and theories on aging and memory.

### Memory and Aging

Episodic memory, defined as the ability to recall specific events and experiences, is known to decline with age. This decline is particularly evident in tasks that require mnemonic discrimination—the ability to distinguish between similar memories. In our study, we employed the MST to measure this ability, a task known to tax hippocampal function and based on modern theories of role of hippocampus in memory (Stark et al., 2019). Consistent with previous research (Yassa et al., 2011; Stark et al., 2013, 2019; Adams et al., 2022; Vanderlip et al., 2024a) we observed significant age-related impairments in mnemonic discrimination performance. This finding supports the idea that distinguishing between similar items becomes more challenging with age.

While recognition memory on the MST is known to remain stable across ages in previous studies, our findings diverged slightly in this respect - we observed lower recognition memory in older adults. This effect, however, was largely driven by the younger adults’ better-than-normative performance on the MST recognition task. Younger participants scored about 8% better than the established norm (MEAN_20-39, Stark 2013_ = 78.6% vs MEAN _current study_ = 85.3%), while the older adult’s performance was reflective of the large-scale normative data (Stark et al., 2013).

Further investigation into episodic memory was conducted using the RAVLT, a widely used list learning paradigm. Our findings aligned to prior work demonstrating age-related impairments in RAVLT (Stark and Stark, 2017; Venkatesh et al., 2020; Dias et al., 2021). Importantly, RAVLT is known to engage hippocampal regions but also interacts with other cognitive processes, including executive function (Bolla-Wilson and Bleecker, 1986; Consonni et al., 2017; Dörr et al., 2023). These results underscore the multidimensional nature of age-related memory decline, which not only affects episodic memory but also interactions with other cognitive systems.

While older adults, on average, perform worse on both the RAVLT and mnemonic discrimination tasks compared to younger individuals, this decline varies widely among individuals. Factors such as education, overall health, and the presence of neurodegenerative diseases like Alzheimer’s can influence the degree of cognitive decline (Salthouse, 2012). The variability in performance underscores the utility of individual differences in memory measures (both RAVLT and MST) for detecting subtle deficits in healthy aging populations and predicting the onset of mild cognitive impairment (MCI) and Alzheimer’s disease (AD).

### Lower GABA levels in healthy older adults

We found that older adults have lower raw posterior cingulate GABA+ levels compared to younger adults. These findings are consistent with previous animal research showing reduced GABA levels in rat’s cingulate (Ling et al., 2005). It contributes to growing literature showing age-related decline in raw GABA levels in human older adults compared to younger adults across several brain regions, including the occipital, auditory, frontal, parietal and motor cortex (Gao et al., 2013; Chamberlain et al., 2019; Zuppichini et al., 2024). While age-related GABA reductions are widespread, they are not uniform. For instance, age-related declines are more pronounced in frontal regions than posterior ones (Porges et al., 2017).

The posterior cingulate cortex (PCC) is of particular interest due to its role in memory and its association with AD. Previous studies have highlighted the importance of the PCC in the early stages of AD, where dysfunction, including hypermetabolism and decreased connectivity, can be observed even before significant cognitive decline becomes evident (Buckner et al., 2005, 2009; Zhou et al., 2008). Reductions in GABA levels in the PCC have also been found in individuals with MCI (Fu et al., 2023). PCC is also a component of the default mode network (DMN), particularly vulnerable to the neurodegenerative processes of Alzheimer’s, and its early involvement is even considered a hallmark of AD progression (Zhou et al., 2010). As such, understanding GABA alterations in the PCC during healthy aging may provide critical insights into the pathophysiology of AD and highlight potential targets for early diagnostic biomarkers and therapeutic interventions.

Additionally, GABA+ level declines in aging might be mediated by changes in GABAergic activity at synapses, specific reduction of inhibitory GABA neurons, or a general age-related decline in total grey matter volume. Our fully tissue-composition corrected GABA estimates did not show an age-related decline in GABA levels (consistent with prior reports (Maes et al., 2018)) suggesting that observed reductions in GABA+ levels could at least be partly attributed to tissue composition changes. However, even with tissue correction, we found that GABA+ levels in the PCC remained significantly correlated with individual differences in episodic memory performance, suggesting that GABA decline is a crucial aspect of aging-related memory deficits regardless of the major underlying reason behind this decline.

### GABA and Memory

Individual differences in GABA levels have been associated with cognition in a region-domain specific manner in older adults. For example, GABA levels in the motor cortex are linked with sensorimotor performance (Cassady et al., 2019), visual GABA levels with visual perception (Lalwani et al., 2021) and auditory levels with auditory function (Lalwani et al., 2019; Dobri and Ross, 2021). We thus hypothesized and found that posterior cingulate GABA levels are positively associated with episodic memory scores.

The PCC, as part of the DMN, plays a vital role in memory retrieval, self-referential thinking, and the integration of information across brain regions. Functional imaging shows increased PCC activity during tasks involving past recollection (Spreng et al., 2009) while reduced PCC activity correlates with memory impairments and cognitive decline (Mormino et al., 2012) in AD. Studies suggest that age-related declines in episodic memory and in AD are linked not only to hippocampal changes but also to disruptions in broader brain networks, including regions involved in executive control and attention, like PCC (Greicius et al., 2004; Dennis and Peterson, 2012; Dennis et al., 2014) indicating a broader role of PCC in episodic memory processing. Moreover, studies have shown that there are connections between the PCC and the hippocampus, which help regulate the flow of information and prevent overactivity that when compromised could impair memory function (Kubota et al., 2013). Indeed, reduced connectivity between medial temporal lobe and PCC is associated with episodic memory retrieval problems (Vanneste et al., 2021). These interactions might be mediated by GABAergic interneurons, which can modulate the excitability of both regions and facilitate the proper integration of memory-related information. These might explain the observed relation of PCC GABA+ levels in age-related decline in episodic memory.

Interestingly, our analysis revealed that the relationship between RAVLT scores and GABA+ levels was not significant after controlling for tissue composition, while the association between GABA+ and mnemonic discrimination was statistically significant only after controlling for tissue composition. We hypothesize that this difference arises because RAVLT delayed recall is not a purely episodic memory measure; it is a more composite task influenced by factors such as strategy, attention, and other cognitive abilities (Consonni et al., 2017; Dörr et al., 2023). As a result, global age-related brain changes, like cortical thinning, likely contribute to RAVLT performance alongside GABA levels. Supporting this, previous research has linked atrophy to delayed recall performance on the (Ahmed et al., 2018) and demonstrated that RAVLT scores can be predicted from grey matter estimates using machine learning (Moradi et al., 2017). In contrast, the relationship between GABA+ and mnemonic discrimination may be more directly linked to GABAergic changes in the PCC, as this task specifically taxes hippocampal function (Stark et al., 2019) and relies more on the PCC-hippocampus interaction, which is critical for discrete memory processes.

### Limitations & Future Research

A significant limitation of our study is the lack of participants diagnosed with MCI or AD. Future research should investigate how GABA levels in the PCC interact with amyloid-beta (Aβ) plaques and other AD biomarkers such as p-tau and myo-inositol. This would help clarify the role of GABAergic dysfunction in AD progression and whether GABA+ levels can serve as an early biomarker for the disease. Previous research has shown that GABAergic dysfunction in AD is associated with memory deficits and neuronal hyperexcitability, which may accelerate cognitive decline (Palop and Mucke, 2010; Vossel et al., 2013; Corriveau-Lecavalier et al., 2024). Therefore, targeting GABAergic dysfunction early in the disease process may help slow progression and preserve cognitive function (Putcha et al., 2011). Additionally, future research should explore whether GABA mediates age-and AD-related connectivity changes in the DMN (Greicius et al., 2004; Damoiseaux et al., 2008; Hu et al., 2013; Goelman et al., 2024) and how alterations in hippocampal GABA levels together contribute to episodic memory deficits.

## Conclusion

In conclusion, our study highlights the link between individual differences in age-related PCC GABA decline and episodic memory deficits. These findings emphasize the role of GABAergic changes in memory decline and suggest that targeting PCC GABA levels could help slow age-related memory loss.

